# The Emotional Glue of the Senses: Affective Certainty and Congruency in the Rubber Hand Illusion

**DOI:** 10.1101/401422

**Authors:** Maria Laura Filippetti, Louise P. Kirsch, Laura Crucianelli, Aikaterini Fotopoulou

## Abstract

Our sense of body ownership relies on integrating different sensations according to their temporal and spatial congruency. Nevertheless, there is ongoing controversy about the role of *affective congruency* during multisensory integration, i.e. whether the stimuli to be perceived by the different sensory channels are congruent or incongruent in terms of their affective quality. In the present study, we applied a widely used multisensory integration paradigm, the Rubber Hand Illusion, to investigate the role of affective, top-down aspects of sensory congruency between visual and tactile modalities in the sense of body ownership. In Experiment 1 (N = 36), we touched participants with either soft or rough fabrics in their unseen hand, while they watched a rubber hand been touched synchronously with the same fabric or with a ‘hidden’ fabric of ‘uncertain roughness’. In Experiment 2 (N = 50), we used the same paradigm as in Experiment 1, but replaced the ‘uncertainty’ condition with an ‘incongruent’ one, in which participants saw the rubber hand being touched with a fabric of incongruent roughness and hence opposite valence. We found that certainty (Experiment 1) and congruency (Experiment 2) between the felt and vicariously perceived tactile affectivity led to higher subjective embodiment compared to uncertainty and incongruency, respectively, irrespective of any valence effect. Our results suggest that congruency in the affective top-down aspects of sensory stimulation is important to the multisensory integration process leading to embodiment, over and above temporal and spatial properties.

## Introduction

The sense of body ownership, defined as the feeling that one’s body belongs to oneself, is considered a fundamental aspect of self-awareness (Gallagher, 2000). A long experimental tradition has attempted to study this facet of bodily self-consciousness by focusing on the self-attribution of visually-presented body parts (e.g. Frassinetti, Maini, Romualdi, Galante, & Avanzi, 2008; Saxe, Jamal, & Powell, 2006). About 20 years ago, a different research tradition emerged that investigates feelings of body ownership in relation to processes of multisensory integration in personal and peripersonal space (Holmes & Spence, 2005; Stein & Standford, 2008). For example, in the Rubber Hand Illusion (RHI; Botvinick & Cohen, 1998), a rubber-hand viewed in peripersonal space (i.e. space near to the body) is experienced as part of one’s body if the subject’s own hand is synchronously touched out of view.

There are now hundreds of studies on the RHI, and related psychophysical or, virtual reality paradigms (see Kilteni et al., 2015 for a review). These investigations have attempted to specify the principles by which multisensory integration happens and the role of different sensory and motor modalities in this integration. Moreover, a number of studies have focused on the role of various ‘amodal’ properties, such as temporal synchrony and spatial congruency in the integration process. For instance, the subject’s hand posture, spatial position and orientation, the equivalent spatial characteristics of the rubber hand, and the relation of both to the midline of the body have been found to modulate the RHI (Preston 2013; Costantini & Haggard, 2007; Graziano, Cook, & Taylor, 2000; Tsakiris & Haggard, 2005). Importantly, top-down processes, defined as information processing that is guided by an individual’s higher-level knowledge and expectations, have also been found to mediate the RHI and related illusions. For example, the RHI is abolished when participants watch a neutral object instead of a hand being stroked synchronously with their own hand (Graziano et al., 2000; Tsakiris & Haggard, 2005; Tsakiris, Costantini, and Haggard, 2008), while subjects can experience a RHI even when touch on the rubber hand is merely expected rather than experienced (Ferri et al., 2013). These findings suggest that the RHI is mediated by both bottom-up processes of multisensory integration and top–down expectations (Tsakiris, 2010; Apps & Tsakiris, 2014).

Moreover, recent computational models have formalised these combined influences. Predictive coding (Zeller et al., 2015) and Bayesian causal inference (Samad, Chung and Shams, 2015) models suggest that the illusion relies on Bayes-optimal inferences about the causes of certain sensations (in this case the inferred common cause is ‘my body’), based on prior knowledge and the spatio-temporal congruency conveyed by these sensations during the RHI.

More recently, the integration of interoceptive modalities, defined as the sensory channels conveying the physiological state of the body, have been found to play a role in the RHI alongside exteroceptive modalities (e.g. Panagiotopoulou et al., 2017; Suzuki et al., 2013; Crucianelli, Metcalf, Fotopoulou, & Jenkinson, 2013; Lloyd, Gillis, Lewis, & Farrell, 2013; van Stralen et al., 2014; Capelari et al., 2009). For example, tactile stimuli that are either painful (Capelari et al., 2009) or pleasant (van Stralen et al., 2014; Crucianelli et al., 2013) in nature lead to strengthened RHI effects. However, it remains unclear whether these effects are caused by the bottom-up, interoceptive specificity and resulting affective value of the felt touch on the real hand, or instead by the congruency between this felt affectivity and the vicariously perceived affectivity of the seen rubber hand. Indeed, it is well recognised that both unpleasant (painful) touch and affective (pleasant) touch can be perceived vicariously when seen on other bodies, i.e. the brain simulates the experience of touch, including its affective aspects (Morrison, Löken, and Olausson, 2010; Thomas, Press, & Haggard, 2006; Keysers et al., 2004). When it comes to the perception of our own body, these two routes, e.g. direct and ‘vicarious’ perception of affective touch, concern the same object and hence require cross-modal integration. However, it is unclear whether incongruences between the felt and seen tactile affectivity can disrupt the multisensory integration processes that lead to body ownership, over and above any effects of temporal, or spatial synchrony.

Indeed, this research tradition has paid little attention to the role of *affective congruency* during multisensory integration. Affective congruency in such paradigms refers to whether the stimuli to be perceived by the different sensory channels, such as the felt touch on the participant’s own hand and visual perception of touch on a rubber hand, are congruent or incongruent in terms of their affective quality. For example, touch on the real hand may be pleasant in valence and/or high in arousal, while touch on the rubber hand may be unpleasant and/or low in arousal, and vice versa. Recent theoretical proposals suggest that affective contingency in embodied interactions may be critical for the formation of the sense of self (Ciaunica & Fotopoulou, 2016; Fotopoulou & Tsakiris, 2017). Indeed, a plethora of developmental studies has highlighted the formative role of affective congruency or contingency in early embodied (multisensory) interactions with caregivers (Reddy et al., 2007; Gergeley & Watson, 1999; Watson, 1994; Bahrick & Watson, 1985). Nevertheless, while the role of temporal and spatial congruency between the felt and seen modalities has been extensively studied in body ownership research, to our knowledge only two studies have manipulated affective congruency vs. incongruency in illusions of body ownership. Moreover, as we explain below, both should be regarded as preliminary given their small samples and incomplete designs.

Specifically, Kammers and colleagues (2011) tested nine healthy participants and used a thermode to induce temperature changes in the ‘real’ hand during a classic RHI procedure, without any corresponding visual manipulations of the rubber hand. They found no differences in RHI strength between neutral and ‘painful’ conditions, i.e. in conditions where cooling or heating of the real hand reached noxious levels. However, they did observe that cooling of the real hand produced greater proprioceptive drift (but no change in subjective ratings) compared to warming of the real hand. These results are hard to evaluate given the small sample, as well as the fact that the paradigm did not involve manipulations of the perceived temperature of the seen, rubber hand that could have been perceived ‘vicariously’. Hence, it is unclear whether the observed effects relate to the provision of additional interoceptive information on the real hand, the increased arousal caused by such stimulation, or the affective incongruency between the real hand (e.g. experienced as warm or cold) and the observed rubber hand (experienced as non-informative with regards to temperature). To our knowledge the only study to include affectivity manipulations on the rubber hand in relation to the real hand, used the RHI paradigm to investigate whether participants’ perception of soft, pleasant versus rough, unpleasant tactile stimulation on their own hidden arm was modulated by simultaneously watching the rubber hand being touched by either the same or an affectively incongruent material (i.e. soft and rough fabric; Schütz-Bosbach et al., 2009). The perceived pleasantness of these fabrics was manipulated based on their softness/roughness, which in turn relates to the perception of the spatial density of the features on a material’s surface (for an overview see Lederman & Klatzky, 2004). However, spatial density is not identical to perceived softness, which shows interindividual variability (see e.g., Bergmann, Tiest & Kappers, 2007) and it is thus a combination of sensory perception and top-down interpretation. Their results did not show a general effect of affective congruency on the RHI, nor did they find a valence effect between the incongruent conditions, e.g. soft touch on the real hand leading to greater embodiment of a rubber hand that was touched by a rough fabric, or vice versa. However, their experiment involved a between-subject design with only 15 participants per group, which may have been underpowered given the multifactorial design, the noise and individual differences involved in such paradigms.

In the present study, we investigated the role of *affective congruency* between modalities in the sense of body ownership over two separate within-subjects RHI experiments (Experiment 1: N = 36; Experiment 2: N = 50). We manipulated for the first time (1) the degree of certainty of the seen tactile affectivity in relation to the felt tactile affectivity and (2) the congruency between seen and felt tactile affectivity. Specifically, in a first experiment, we manipulated the pleasantness of fabrics touching the real hand (based on their relative softness), whilst the rubber hand was exposed to temporally and spatially congruent touch that was either also affectively congruent or affectively ambiguous. That is, participants’ real hidden arm was stroked by a pleasant (i.e. synthetic wool) or non-pleasant (i.e. Velcro) fabric while they watched a rubber hand receiving a synchronous tactile stimulation by the same congruent fabric or by a ‘hidden’ fabric. We asked the question of whether *affective certainty (vs. uncertainty)* can affect changes in embodiment, over and above the effects of temporal synchrony (vs. asynchrony). Experiment 2 used an identical paradigm to the one described in Experiment 1 but replaced the emotionally uncertain experience of the hidden fabric on the rubber hand with an emotionally incongruent fabric. That is, participants watched a rubber hand being touched by a pleasant (i.e. synthetic wool) or non-pleasant (i.e. Velcro) fabric while they received on their real hidden arm a synchronous tactile stimulation by the same congruent fabric or by the incongruent fabric. We asked the question of whether *affective congruency (vs. incongruency)* can create changes in embodiment, over and above the effects of temporal synchrony (vs. asynchrony). Indeed, in both experiments, we also included a control asynchronous condition, in which both the participant’s hand and the rubber hand were stroked using a pleasant fabric, but the stimulation was offset. Further, in both experiments, we measured the RHI in two ways: 1) by asking participants to localise their unseen hand (i.e. proprioceptive drift), and 2) by a questionnaire investigating the subjective change in body ownership and awareness (i.e. embodiment questionnaire; Longo et al., 2008). Additionally, we measured the perceived pleasantness of the felt touch to investigate the relation between changes in embodiment and the perceived tactile affectivity.

We hypothesised that if affective congruency between the ‘felt’ and ‘seen’ touch is important in embodiment, the RHI effects will be weaker in conditions in which (1) there is uncertainty about the affective congruency of visual and tactile stimulations, despite their temporal and spatial congruency; (2) visual and tactile stimulations are affectively incongruent, despite their temporal and spatial congruency. If on the other hand, affective valence rather than congruency is important to the multisensory integration process, then we should observe a valence effect, i.e. the illusion may be enhanced by pleasant touch versus non-pleasant touch, over and above any effect of certainty or congruency. Additionally, we also expected predictable affectivity (i.e. certain and congruent seen fabrics) to lead to greater pleasantness ratings than unpredictable affectivity (i.e. incongruent seen fabrics). Alternatively, if the ‘valence’ of the felt affectivity is uniquely important during multisensory integration, then we should observe valence effects in the second experiment (i.e. pleasant fabrics applied to the hand should lead to greater pleasantness ratings during the RHI than non-pleasant fabrics applied to the real hand, irrespective of the affective congruency manipulations).

## Methods

### Participants

Ninety-four female participants were recruited to take part in the study. Eight participants were excluded due to the following reasons: (1) experimental error and hence only partial testing (Experiment 1: N = 2), (2) rating the Velcro fabric as more pleasant than the synthetic wool at the baseline tactile test (Experiment 1: N = 2; Experiment 2: N = 3) and (3) failing to follow the instructions during the induction of the RHI (Experiment 2: N = 1). Thus, the final sample consisted of 86 participants (Experiment 1: N = 36, mean age = 25.11 years, SD = 9.78 years; Experiment 2: N = 50, mean age = 21.25 years, SD = 3.23 years). Institutional ethical approval was obtained and the experiment was conducted in accordance with the Declaration of Helsinki. Informed written consent was obtained from each participant prior to the experiment.

### Materials

The participant’s left hand and the rubber hand were positioned in a black wooden box measuring 34 × 65 × 44 cm (Crucianelli et al., 2013), which hides the participant’s own arm from view but allows the rubber hand to be viewed. Hence, during the RHI participants were not able to see the fabric that was stroking their own real arm, but only to perceive it at the tactile level. The stroking was done by the experimenter using a black plastic stick, to which the respective fabric was attached. The size and colour of the fabrics were matched to avoid perceptual confounds. The choice of the two fabrics was made on the basis of a pilot experiment, whereby 12 materials of different degrees of pleasantness (cotton wool ball, soft sponge, Velcro hook, Velcro hairy loops, hair doughnut, faux fur, sandpaper, linen fabric, wool yarn, latex, cotton fabric, synthetic wool yarn – please see Figure 1S Supplementary Material) were contrasted in a visual only, tactile only, and visuo-tactile conditions. Based on these data, we chose the most pleasant and least pleasant (hereafter referred to as non-pleasant) fabrics with similar visual appearance (i.e. size and colour), as rated by 12 participants.

In Experiment 1, we contrasted the two congruent conditions (felt and seen touch was either pleasant or non-pleasant) against two conditions of uncertain affectivity (see Figure 1. A). In the latter two conditions, the participant’s hand was stroked using either the pleasant or the non-pleasant fabric, while the fabric stroking the rubber hand was hidden from participant’s view. To hide the fabric, the experimenter used a stick with a cardboard attached, which prevented the participants from viewing just the tip of the stick. Thus, participants could clearly see where and how fast the rubber hand was touched, but they could not see the precise fabric that was touching it. In Experiment 2, we replaced the manipulation of the certainty of affectivity with the manipulation of the congruency of affectivity (see Figure 1. B). Thus, in this experiment participant’s hand was stroked either congruently in two conditions as above (felt and seen touch was either pleasant or non-pleasant), or incongruently in two, new conditions (felt touch was pleasant and seen touch was non-pleasant and vice versa). Finally, in both experiments, in the asynchronous condition, both the participant’s hand and the rubber hand were stroked using the pleasant fabric, but the stimulations were offset.

**Figure 1.**
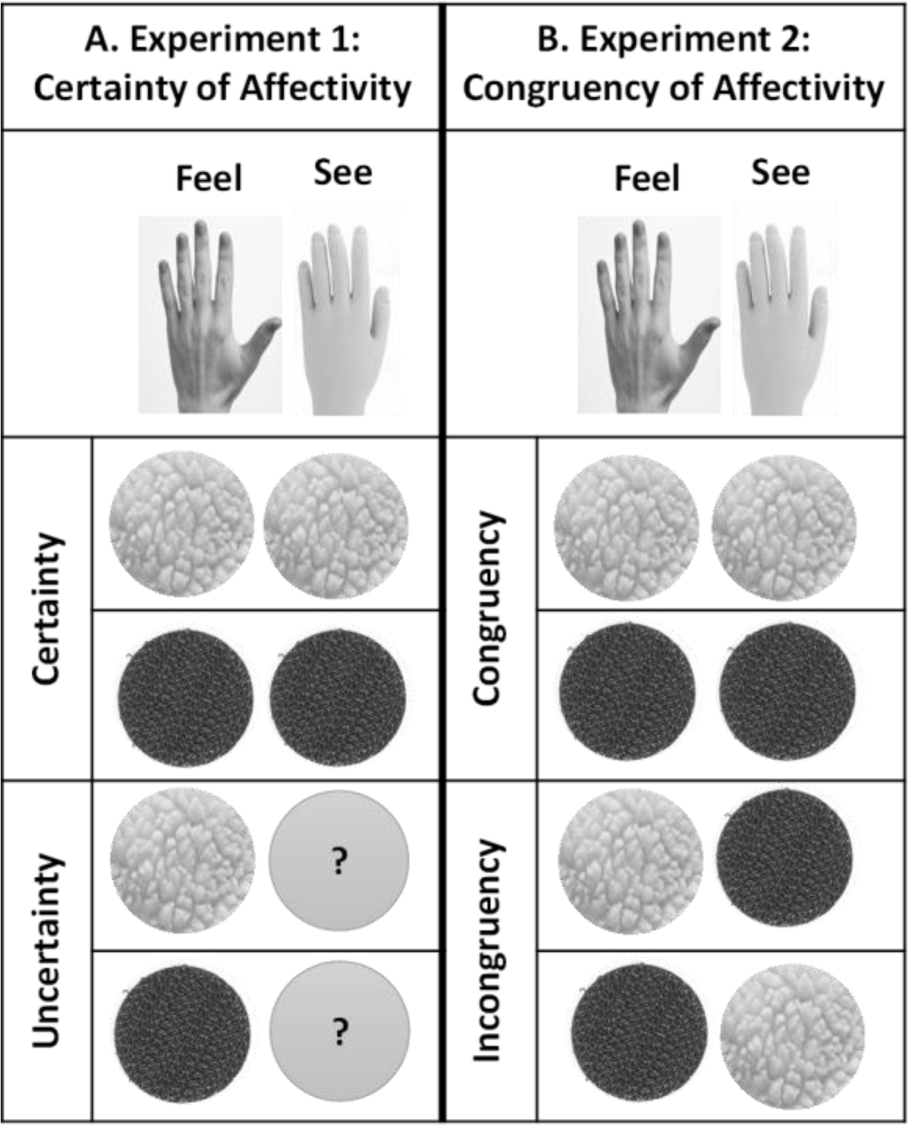
Illustration of the experimental design of Experiments 1 and 2. A. Experiment 1 consisted in a 2×2 within-subject design, with the independent variables Fabric (pleasant vs. non-pleasant) and Certainty of Affectivity on the rubber hand (congruent vs hidden). B. In Experiment 2, we contrasted the independent variables Fabric (pleasant vs. non-pleasant) and Congruency of Affectivity on the rubber hand (congruent vs incongruent). In both experiments, an additional asynchronous condition (not illustrated here) was used to control for the strength of the RHI. This condition used the pleasant fabric (synthetic wool) on both the real and rubber hands and was contrasted against the pleasant congruent condition.

### Experimental setup and design

As illustrated in Figure 1, both experiments followed a within-participant repeated measures design. The independent variables of Experiment 1 were Fabric (pleasant vs. non-pleasant) and Certainty of Affectivity (congruent certain vs. hidden/uncertain fabric). In this experiment, participants left hand was stroked using a pleasant (synthetic wool) or non-pleasant (Velcro) fabric, whilst they watched a rubber hand being stroked by the same material (i.e., pleasant or non-pleasant conditions; congruent certain condition), or by a hidden fabric (hidden/uncertain condition). An additional asynchronous condition was included to control for the strength of the illusion using the pleasant fabric only (asynchronous pleasant congruent condition). The order of conditions was randomised across participants. Our dependent variables were: (1) An embodiment questionnaire (Longo et al., 2008), to provide a measure of embodiment over the rubber hand (18 statements, 7-point Likert-type scale; −3, strongly disagree; +3, strongly agree). In each condition, the questionnaire was administered pre-(i.e., embodiment due to the visual capture effect; Martinaud et al., 2017) and post-stroking. We calculated their difference to obtain a measure of subjective embodiment due to visuo-tactile integration (post – pre measures). We computed the ‘embodiment score’, obtained by averaging the scores of ownership, felt location and felt agency together (Longo et al., 2008). (2) A proprioceptive drift measure, that is the degree to which the stimulated hand is perceived to be closer to the rubber hand. In each condition, we first subtracted the value corresponding to the actual position of the participant’s index finger from the value corresponding to the felt position, before (“pre” value) and after (“post” value) stroking. Then their difference (post – pre) was calculated to obtain a measure of proprioceptive drift due to rubber hand illusion induction. (3) A subjective valence rating of stroking per condition (participants are asked to indicate the extent of their agreement or disagreement with three statements: “How pleasant was it on your skin”, “How unpleasant was it on your skin”, and “How arousing was it on your skin” on a VAS with two anchors (0, not at all; +100, extremely), to measure the perceived affectivity of the various conditions.

In Experiment 2, while we kept the independent variables Fabric (pleasant vs. non-pleasant) identical, we replaced the certainty/uncertainty of the seen fabric with a congruent vs. incongruent manipulation (Figure 1.B). That is, participants’ left hand was stroked using a pleasant (synthetic wool) or non-pleasant (Velcro) fabric, whilst they watched a rubber hand being stroked by the same material (i.e., pleasant or non-pleasant conditions) used to stroke their own hand (congruent condition), or by an incongruent fabric (incongruent condition).

In Experiment 1 we used a box frame whereby a realistic left rubber hand was situated 25 cm to the right of the participant’s own left hand. Given recent evidence, we felt that this distance may reduce proprioceptive drift effects and hence in Experiment 2 we used a different wooden box, where the distance between the rubber hand and the real left hand was 17 cm (Preston, 2013; Lloyd, 2007). Aside from this difference, all the other experimental setup procedures were kept identical between the two experiments.

### Experimental procedure

The experimenter marked the stroking skin area measuring 9cm long × 4cm wide with a washable marker on the hairy skin of participants’ left forearm (wrist crease to elbow, McGlone et al., 2012; Crucianelli et al., 2013). Next, the participant’s left arm and the rubber hand were positioned in the black wooden box. Before the stroking stimulation, the rubber hand was hidden to take a baseline measure of pointing response (for the proprioceptive drift measure); the experimenter ran her finger along the length of the ridge of the box until the participant would tell the experimenter to stop when her finger was directly over the location of the participant’s hidden index finger (Lloyd et al., 2013). Then, before the experimental procedure started, participants were asked to complete a baseline tactile test. The experimenter then delivered tactile stimulation on the participant’s hidden forearm with either the synthetic wool (pleasant fabric) or Velcro (non-pleasant fabric). The stimulation was done once and participants were then asked to complete three statements of valence ratings (“How pleasant was the touch”, “How unpleasant was the touch”, and “How arousing was the touch”: on a 0 to 100 VAS, with 0, not at all; +100, extremely).

Then, the rubber hand was uncovered and the experimenter asked the participant to look at the rubber hand without moving his/her own hidden hand for 15 s. The participant was then required to fill the (pre) embodiment questionnaire. This measure of subjective visual capture was collected before each experimental condition and was then contrasted against the (post) embodiment questionnaire collected after the visuo-tactile induction. The experimenter then sat in front of the participant and manually delivered stimulation to the visible rubber hand and the participant’s hidden forearm. Participants were stimulated on their left forearm, with a stroking velocity of 3 cm/s, matching in both space and time. In the Pleasant Congruent and Non-pleasant Congruent conditions, the participant’s hand and the rubber hand were stroked simultaneously in the same anatomical location using either the pleasant (synthetic wool) or the non-pleasant (Velcro) fabrics.

In all conditions, participants were instructed to keep their own left hand still and carefully observe the rubber hand. Each stroking stimulation lasted for 2 minutes. Immediately following the stroking stimulation, the experimenter recorded the proprioceptive drift, (post) embodiment measures and VAS valence ratings as explained in the Experimental setup. Prior to commencing the next condition, participants were given a rest period during which they were instructed to freely move and take the hand out of the box.

After the experimental procedure was completed, participants underwent a baseline visual test, similar to the baseline tactile test described above. Here, they were asked to watch the rubber hand, whilst the experimenter delivered tactile stimulation with either pleasant or non-pleasant fabric on the artificial hand itself. As in the baseline tactile test, participants completed the three statements of valence ratings (“How pleasant did you think the touch was”, “How unpleasant did you think the touch was”, and “How arousing did you think the touch was”, on a 0 to 100 VAS, with 0, not at all; +100, extremely). As in Schütz-Bosbach et al (2009), these tactile and visual ratings were used to check whether participants could discriminate between the two fabrics at the tactile level and whether they could associate the seen fabrics with the felt tactile stimulation.

## Results

### Embodiment questionnaire

The change in embodiment across experimental conditions (post-pre of the average of scores from the embodiment questionnaire) was analysed using repeated measure non-parametric analyses, as the data were non-normally distributed. A Wilcoxon signed rank test revealed a main effect of Stroking, with synchronous stroking (Experiment 1: Md = 0.94; Experiment 2: Md = 1.25) producing significantly higher embodiment scores than asynchronous stroking (Experiment 1: Md = 0.19; Experiment 2: Md = 0.00), (Figure 2 - Experiment 1: Z = 2.839, *p* = 0.005, r = 0.47; Experiment 2: Z = 5.053, *p* < 0.001, r = 0.71). We also performed the Wilcoxon signed rank test on the main effect of Certainty of Affectivity of the fabric (Experiment 1), which revealed that participants embodied the rubber hand to a significantly greater extent when the stroking material was congruent (Md = 0.68), compared to when it was hidden (Md = 0.22), (Z = −2.826, *p* = 0.005, r = 0.24). An identical non-parametric analysis was performed for Experiment 2, in relation to the main effect of Congruency of Affectivity of the fabric, which revealed that participants embodied the rubber hand to a significantly greater extent when the stroking material was congruent (Md = 1.53), compared to when it was incongruent (Md = 0.69) (Z = −4.403, *p* < 0.001, r = 0.62). In both experiments, the main effect of Fabric on embodiment was not significant (Figure 2), suggesting that participants embodied the rubber hand irrespective to whether the stroking material was pleasant (Experiment 1: Md = 0.75; Experiment 2: Md = 0.81) or non-pleasant (Experiment 1: Md = 0.31; Experiment 2: Md = 1.13), (Experiment 1: Z = −0.631, *p* = 0.524, r = 0.11; Experiment 2: Z = 0.877, *p* = 0.380, r = 0.12). The interactions between Certainty of Affectivity and Fabric (Experiment 1) and Congruency of Affectivity and Fabric (Experiment 2) were analysed by calculating the difference between the experimental manipulation (hidden or incongruent fabric, depending on the experiment) and congruent fabric scores in the pleasant and non-pleasant conditions separately, and subsequently using a Wilcoxon signed rank test to compare these two differential scores. This analysis revealed no significant interactions (Experiment 1: Z = 0.074, *p* = 0.941, r = 0.01; Experiment 2: Z = −1.249, *p* = 0.212, r = 0.03). In sum, our results reveal the typical effect of synchronicity on the subjective experience of the RHI (i.e. greater feelings of RH ownership following synchronous than asynchronous stimulation). We also show for the first time that the subjective experience of the illusion is influenced by affective certainty and congruency (i.e. greater feelings of RH ownership following affective certain and congruent than uncertain or incongruent stimulation). However, no fabric nor interaction effect was observed in either experiment.

**Figure 2.**
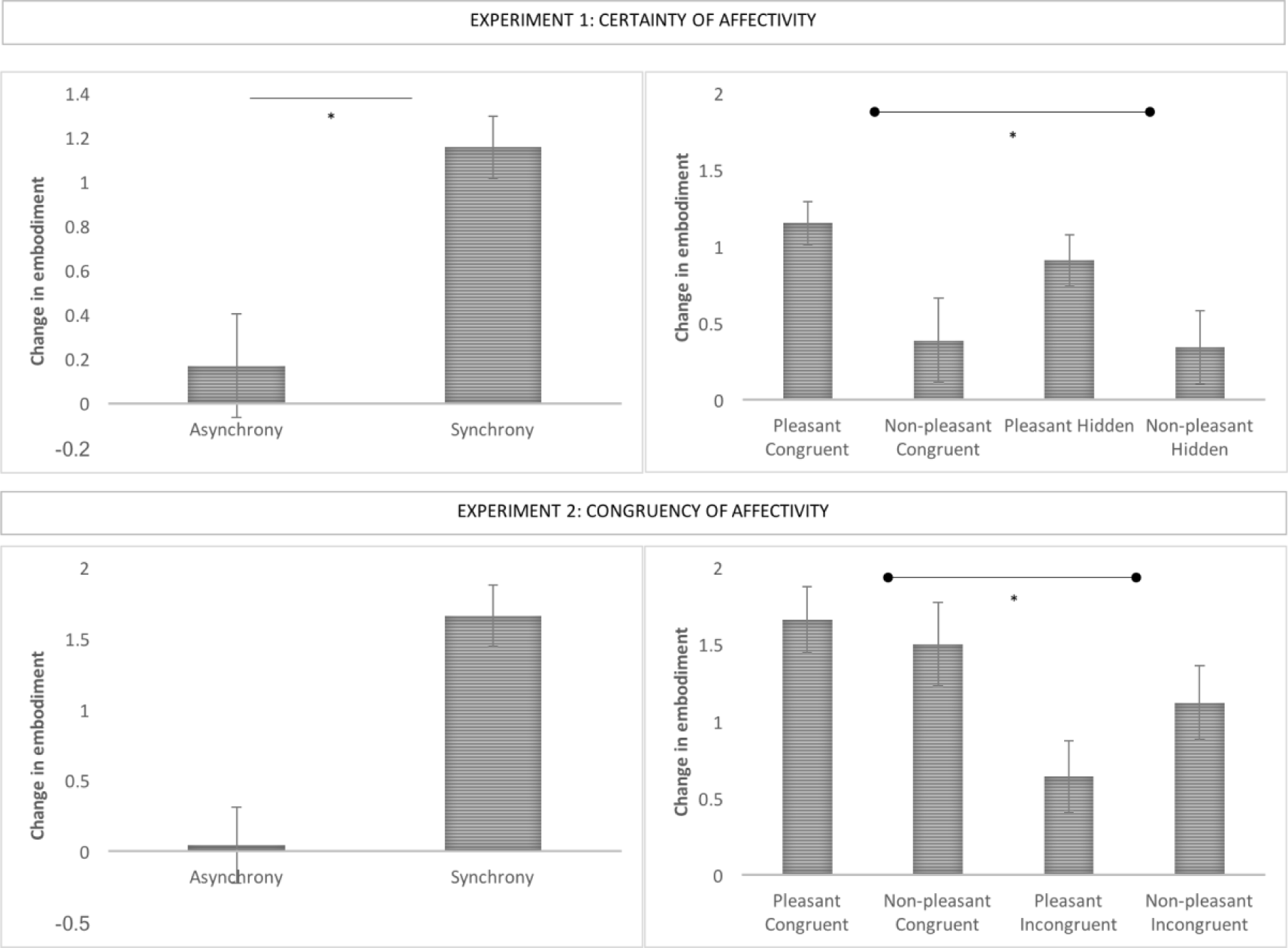
Change in embodiment as measured by subjective reports (embodiment questionnaire) between the synchronous and asynchronous conditions (left side of the figure), and across experimental conditions (right side of the figure). The graphs on the top of the image shows mean differences in Experiment 1, whereas the bottom of the image shows mean differences in Experiment 2. Means are shown for illustrative purposes. Error bars indicate standard error of the mean. Asterisks denote significant differences (p < 0.05).

### Proprioceptive drift

In Experiment 1, data was normally distributed. Hence, we performed a repeated-measure ANOVA with Fabric (pleasant vs non-pleasant) and Certainty of Affectivity (congruent vs hidden fabric) as within-subject variables. This analysis revealed no significant main effects nor interactions (Figure 3 – Experiment 1): main effect of Fabric, F (1,35) = 0.112, *p* = 0.740, *η*_^*2*^_ = 0.003; main effect of Neutral Affectivity, F (1,35) = 1.198, *p* = 0.281, *η*_^*2*^_ = 0.033; interaction Fabric * Certainty of Affectivity, F (1,35) = 2.251, *p* = 0.142, *η*_^*2*^_ = 0.060. A repeated-measure t-test was performed to investigate the difference in drift towards the rubber hand between asynchronous and synchronous (Pleasant Congruent) conditions, which revealed no significant differences, t(35) = −1.235, *p* = 0.225, *d* = 0.15.

**Figure 3.**
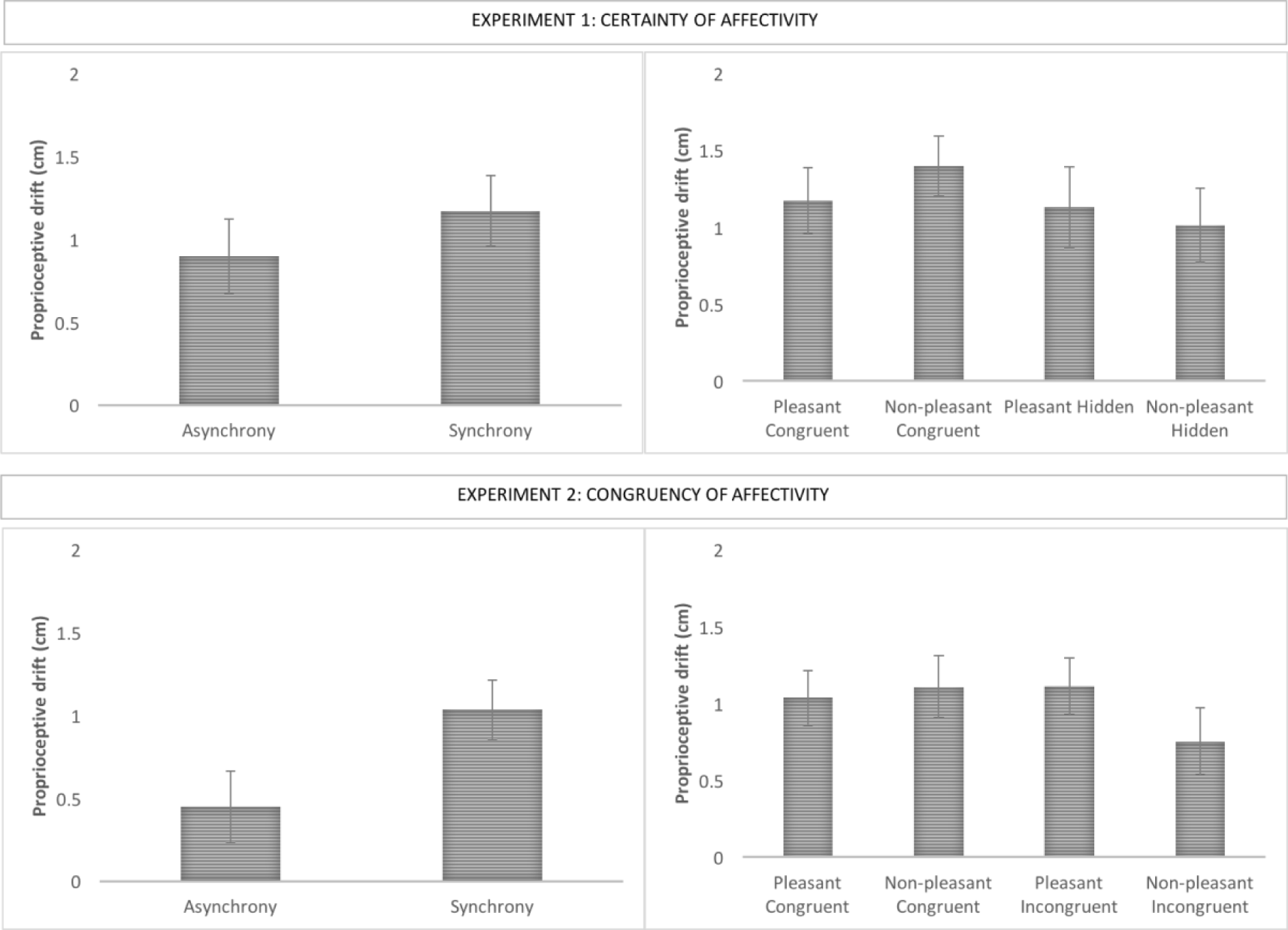
Change in proprioceptive drift between the synchronous and asynchronous conditions (left side of the figure), and across experimental conditions (right side of the figure). The top of the image shows mean differences in Experiment 1, whereas the bottom of the image shows mean differences in Experiment 2. Error bars indicate standard error of the mean. Asterisks denote significant differences (p < 0.05).

Data exploration of Experiment 2 revealed the presence of an outlier (with a score above 2 SD from the mean), which was excluded from the final sample. We performed a repeated-measure ANOVA with Fabric (pleasant vs non-pleasant) and Congruency of Affectivity (congruent vs. incongruent fabric) as within-subject variables. This analysis revealed no significant main effects nor interactions (Figure 3 – Experiment 2): main effect of Fabric, F(1,48) = 1.038, *p* = 0.313, *η*_^*2*^_ = 0.021; main effect of Congruency of Affectivity, F (1,48) = 0.818, *p* = 0.370, *η*_^*2*^_ = 0.017; interaction Fabric * Congruency of Affectivity, F (1,48) = 3.579, *p* = 0.065, *η*_^*2*^_ = 0.069. The comparison between asynchronous and synchronous (pleasant congruent) conditions here revealed a significantly bigger proprioceptive drift towards the rubber hand after synchronous compared to asynchronous condition, t(48) = − 3.255, *p* = 0.002, *d* = 0.37.

In sum, results on proprioceptive drift show a change in self-location as a function of synchronicity only in Experiment 2. However, no main effect (fabrics or affective certainty and congruency) nor interaction was observed in either experiment.

### Valence ratings

To analyse the effects of multisensory stimulation on perceived tactile affectivity, we conducted separate analyses for each of the three relevant measures (pleasantness, unpleasantness, arousal). As the data were non-normal, non-parametric analyses were performed using the Wilcoxon signed rank test.

In Experiment 1, for the pleasantness ratings (Figure 4), we found a main effect of Fabric, in the sense that the pleasant fabric (Md = 83) was rated as more pleasant than the non-pleasant fabric (Md = 35.25) regardless of the experimental condition, Z = −4.737, *p* < 0.001, r = −0.45, and a main effect of Certainty of Affectivity, whereby the congruent condition (Md = 59) was rated more pleasant than the hidden condition (Md = 59.75), regardless of the fabric, Z = −2.199, *p* = 0.028, r = −0.37. We did not find any interaction between Fabric and Certainty of Affectivity, Z = −1.066, *p =* 0.29, r = −0.178. Similarly, for the unpleasantness ratings (Figure 5), we found a main effect of Fabric, with the non-pleasant fabric (Md = 49.25) being rated as more unpleasant than the pleasant fabric (Md = 9), Z = 4.878, *p* < 0.001, r = 0.81. However, we did not find a main effect of Certainty of Affectivity, Z = 1.941, *p* = 0.052, r = −0.32, nor interaction between Fabric and Certainty of Affectivity, Z = 0.786, *p* = 0.43, r =0.13. No main effects nor interactions were found with regard to the arousal ratings (Figure 5) (Fabric: Z= − 1.31, *p* = 0.190, r = −0.22; Certainty of Affectivity: Z = −1.30, *p* = 0.19, r = −0.22). Additionally, there were no differences in pleasantness (Z= −1.521, *p* =0.128, r = − 0.25), unpleasantness (Z =-1.411, *p* = 0.158, r = − 0.49) or arousal (Z = −0.769, *p* = 0.442, r = −0.13) arising from the tactile stimulation between the synchronous and asynchronous conditions.

**Figure 4.**
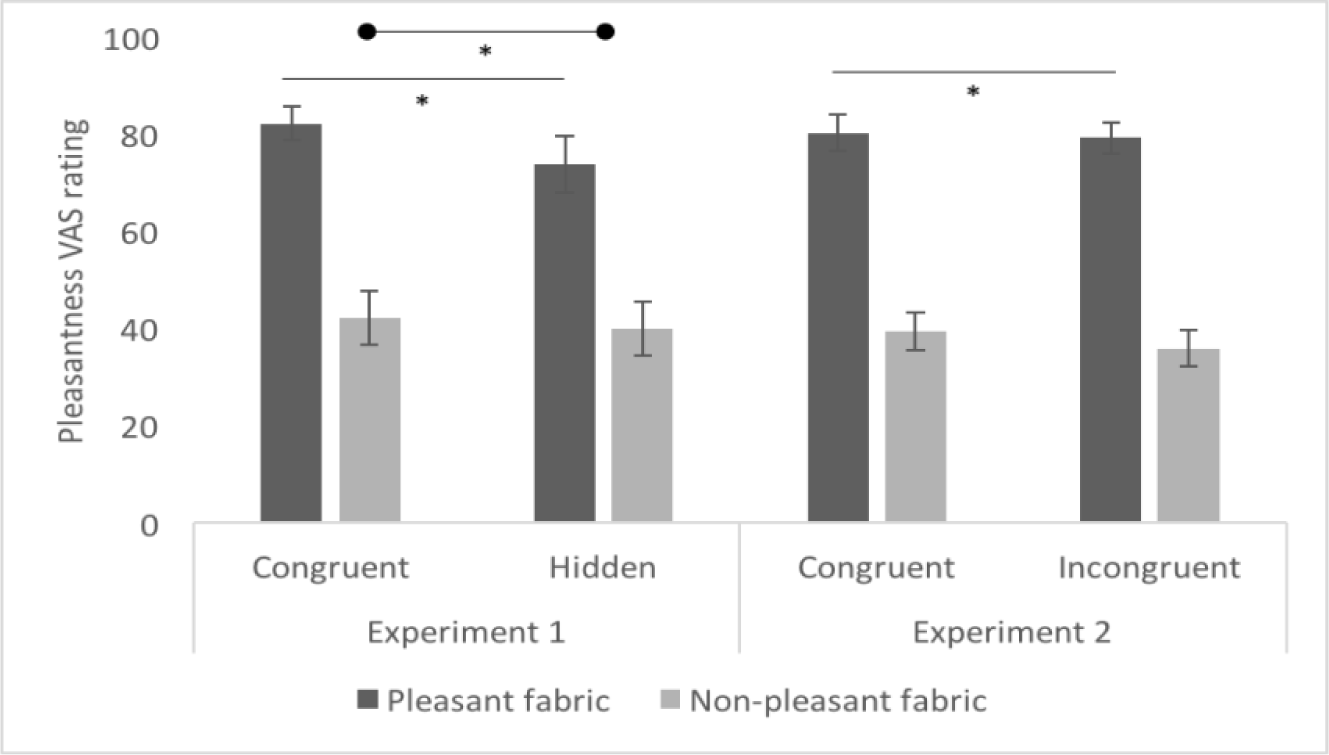
Pleasantness valence ratings post-induction of the RHI in both Experiment 1 and Experiment Means are displayed for illustrative purposes. Error bars indicate standard error of the mean. Asterisks denote significant differences (p < 0.05).

**Figure 5.**
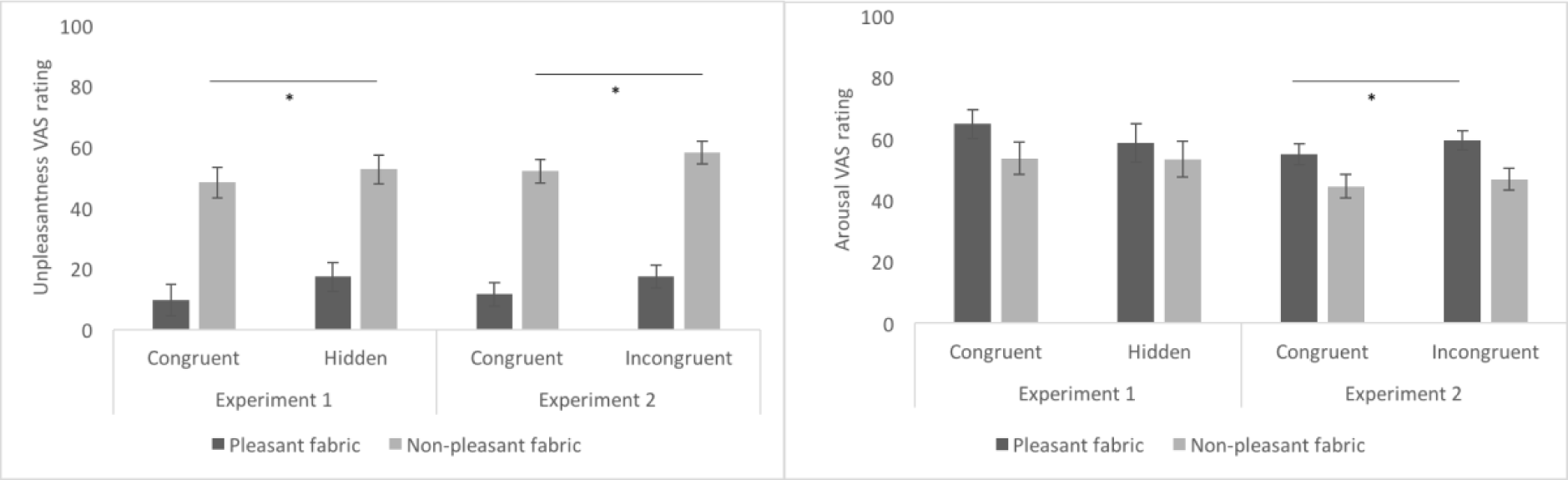
Unpleasantness (left graph) and arousal (right graph) valence ratings post-induction of the RHI in both Experiment 1 and Experiment 2. Means are displayed for illustrative purposes. Error bars indicate standard error of the mean. Asterisks denote significant differences (p < 0.05).

In Experiment 2, for the pleasantness ratings (Figure 4), we found a main effect of Fabric, that is the pleasant fabric (Md = 82.25) was rated as more pleasant than the non-pleasant fabric (Md = 38.25), Z = −5.656, *p* < 0.001, r = −0.80, but no main effect of Congruency of Affectivity, with no differences in pleasantness ratings between congruent (Md = 59.5) and incongruent (Md = 56) visuo-tactile stroking, Z = −0.656, *p =0.512,* r = −0.09. We did not find any interaction between Fabric and Congruency of Affectivity, Z = −1.595, *p* = 0.11, r = 0.23. Similarly, for the unpleasantness ratings (Figure 5), we found a main effect of Fabric, with Velcro (Md = 52.25) being rated as more unpleasant than wool (Md = 5.5), Z = 5.869, *p* < 0.001, r = 0.83, and no main effect of Congruency of Affectivity (Md congruency = 35.5; Md incongruency = 39; Z = 1.694, *p* = 0.090, r = 0.24). No interaction between Fabric and Congruency of Affectivity was found, Z = −0.464, *p* = 0.642, r = −0.07. Participants rated the synthetic wool (Md = 60.5) as more arousing than the Velcro (Figure 5) (Md = 50) (main effect of Fabric, Z = −2.761, *p* = 0.006, r = −0.39), but no main effect of Congruency of Affectivity or interactions between Fabric and Congruency of Affectivity were found on arousal, Z = 1.479, *p* = 0.139, r = 0.21, and Z = −0.717, *p* = 0.473, r = 0.10, respectively. As for Experiment 1, there were no differences in pleasantness (Z = −0.873, *p* = 0.382, r = −0.12), unpleasantness (Z = 1.467, *p* = 0.142, r = 0.21) or arousal (Z = 0.573, *p* = 0.567, r = 0.081) in the tactile stimulation between the synchronous and asynchronous conditions.

In sum, our results reveal that our fabric-based, manipulations of tactile pleasantness were successful in the sense that there were main effects of the felt fabric in both experiments, with the pleasant fabrics being experienced as more pleasant and the non-pleasant fabrics being experienced as more unpleasant overall. We also show that affective certainty has an effect on our valence ratings (i.e. higher perceived pleasantness of the certain condition compared to the hidden condition). However, this effect was not apparent in Experiment 2, in which there were no differences in perceived affectivity between congruent and incongruent conditions. No interaction effect was observed in either experiment. Finally, there were no differences in valence ratings between the synchronous and asynchronous condition in either experiments.

### Tactile and visual discrimination baseline measure

To assess how participants perceived the valence of the two fabrics in unisensory stimulation, i.e. in our touch only and vision only baseline measurements, we analysed their ratings of fabric ‘pleasantness’, ‘unpleasantness’ and ‘arousal’. As the data were non-normal, non-parametric analyses were performed using the Wilcoxon signed rank test. For these manipulation checks, we contrasted the two materials (synthetic wool vs. Velcro) within each valence rating statement (pleasantness, unpleasantness, and arousal). Table 1S (Supplementary Material) shows mean and standard deviation values for illustrative purposes. In Experiment 1, the baseline tactile analysis showed that participants rated the pleasant fabric as more pleasant (Md_Wool_ = 77; Md_Velcro_ = 36.5; Z= −5.030, *p* < 0.001, r = −0.84) and less unpleasant (Md_Wool_ = 8; Md_Velcro_ = 42.5; Z = 4.318, *p* < 0.001, r = 0.72) than the non-pleasant fabric. No differences between materials (Md_Wool_ = 59; Md_Velcro_ = 50) were found for the arousal ratings (Z = −1.237, *p* = 0.216, r = −0.20). The same analysis was performed for the visual test, which showed that participants rated the pleasant fabric touching the rubber hand as more pleasant (Md_Wool_ = 81.5; Md_Velcro_ = 17; Z = 4.966, *p* < 0.001, r = 0.83) and less unpleasant (Md_Wool_ = 9.5; Md_Velcro_ = 74; Z = −5.211, *p* < 0.001, r = −0.87) than the non-pleasant fabric. No differences (Md_Wool_ = 62; Md_Velcro_ = 47) were found for the arousal ratings (Z = 1.411, *p* = 0.158, r = 0.24).

In Experiment 2, the baseline tactile analysis showed that participants rated the synthetic wool as more pleasant (Md_Wool_ = 81; Md_Velcro_ = 38; Z= −5.930, *p* < 0.001, r = −0.84) and less unpleasant (Md_Wool_ = 0; Md_Velcro_ = 54.5; Z = 5.160, *p* < 0.001, r = 0.73) than the Velcro. Additionally, the synthetic wool was rated as more arousing than the Velcro (Md_Wool_ = 57.5; Md_Velcro_ = 38; Z = −2.831, *p* = 0.005, r = 0.33). For the baseline visual test, participants rated the sight of the synthetic wool touching the rubber hand as more pleasant (Md_Wool_ = 87.5; Md_Velcro_ = 21.5; Z = −4.998, *p* < 0.001, r = −0.70) and less unpleasant (Md_Wool_ = 5; Md_Velcro_ = 74; Z = 5.344, *p* < 0.001, r = 0.76) than the Velcro. Similarly to the baseline tactile test, the wool was rated as more arousing than the Velcro (Md_Wool_ = 65; Md_Velcro_ = 32; Z = 2.422, *p* = 0.015, r = 0.34).

In sum, the baseline tactile manipulation check shows that participants rated the pleasant fabric as more pleasant and less unpleasant than the non-pleasant fabric. These results apply to both the tactile and the vision modalities. Additionally, in Experiment 2 participants rated the pleasant fabric as more arousing than the non-pleasant fabric in both the tactile and the visual domain.

## Discussion

Our body is a unique multimodal object that can be perceived through both epistemically private modalities, such as touch and proprioception, but also through vision, as any other observed body in the world. Therefore, the ways in which our brain integrates information about the same body from ‘within’ and from ‘without’ (as when one is observing oneself in the mirror) is important for understanding how we come to build a coherent sense of an embodied self. The aim of this study was to investigate for the first time whether our embodied self is influenced by the *affective* congruency between sensations derived from within the body (felt touch) and from without (seen touch). In particular, we took advantage of the unique set-up of the Rubber Hand Illusion (RHI) to examine whether the affective congruency (versus uncertainty and incongruency) between the ‘felt’ touch on one’s own hand and the vicarious, seen touch on a rubber hand would modulate the experience of rubber hand embodiment and perceived pleasantness, over and above the effects of visuo-tactile synchrony and any valence effects. Based on previous findings about the role of (congruent) affective touch in the RHI (Crucianelli et al., 2013; van Stralen et al., 2014; Lloyd et al., 2014), one could predict that participants would be less willing to embody the rubber hand when their real limb feels pleasant, but a non-pleasant fabric touches the rubber hand and vice versa. Given however the role of affective congruency in the development of the bodily self (Fotopoulou & Tsakiris, 2017), we hypothesised that affective congruency is important to the multisensory integration process leading to embodiment and thus the RHI effects would be weaker when the felt tactile affectivity does not match the vicariously perceived tactile affectivity on the rubber hand, irrespective of valence. Indeed, we found that certainty (Experiment 1) and congruency (Experiment 2) between the perceived and observed tactile affectivity led to higher subjective embodiment compared to uncertainty and incongruency, respectively. This suggests that affective congruency between visual and tactile modalities influences the subjective sense of body ownership. Interestingly, we did not find any interaction effect of congruency and valence in either experiment, suggesting that it is not the positive valence of the felt or seen touch that leads to an increased sense of embodiment but rather the congruency between the felt and seen touch, irrespective of valence.

In a previous study, Capelari and colleagues (2009) induced the RHI using nociceptive stimuli by means of sharp pins that were used to synchronously and asynchronously stroke the participant’s hand and the rubber hand. They found that both tactile painful stimuli and purely tactile stimuli elicited similar changes in body ownership. Here we confirm a similar effect by showing that congruency between felt and seen touch leads to higher subjective embodiment irrespective of its pleasantness, and we also show that incongruence between these stimuli reduces the RHI.

Our findings indicate that affective congruency of visuo-tactile stimulation is the main driver of the illusion, thus suggesting that a matching between ‘felt’ and ‘seen’ tactile affectivity may be a key component to induce a change in the conscious experience of body ownership. This is also in line with developmental studies showing that early multisensory contingent interactions with caregivers are key to the construction of a minimal body awareness (Reddy et al., 2007; Gergeley & Watson, 1999; Watson, 1994; Bahrick & Watson, 1985). As recently put forward by Fotopoulou and Tsakiris (2017) our most minimal sense of body awareness might originate from early proximal tactile interactions with other bodies, which simultaneously carry information about the inner body and the external world. Here, we suggest that *affective* congruency between ‘felt’ and ‘seen’ tactile stimulation could represent another type of amodal property of multisensory integration (Bahrick & Lickliter, 2002). In turn, affective incongruency between different sensory channels might create a form of deafferentiation similar to the exteroceptive *temporal* mismatch seen in multisensory asynchrony (Longo et al., 2008). Future studies should further investigate this hypothesis.

While in both experiments embodiment questionnaire results confirmed that participants experienced the RHI, we found a proprioceptive drift effect as a function of synchronicity only in Experiment 2. Recent studies suggest that the distance used in Experiment 1 between the real and the rubber hand may have been suboptimal for the generation of proprioceptive errors (Preston et al., 2013). Additionally, our affective congruency manipulations did not result in any main effect nor interactions on proprioceptive drift measures in either experiment. Thus, our findings are limited to subjective aspects of body ownership. Indeed, recent studies have shown that subjective embodiment and proprioceptive drift can dissociate (Rohde, Di Luca, & Ernst, 2011) and the latter should not be considered as a behavioural correlate of the subjective experience of the illusion (Abdulkarim & Ehrsson, 2016; Holle et al., 2011).

In both experiments, we also measured the perceived pleasantness of the felt touch to investigate the perceived affectivity of affective congruency between felt and seen touch during the RHI (versus uncertainty and incongruency). In Experiment 1, certainty between seen and felt touch lead to greater pleasantness ratings than uncertainty, as predicted. In our second experiment however, congruency was not perceived as more pleasant than incongruency. It is unlikely that this effect was related to participants’ inability to recognise the presence of congruent vs. incongruent tactile affectivity. The baseline measures taken before and after the RHI indicate that participants discriminate between the two fabrics both at the visual and tactile levels. Additionally, in both experiments we found a main effect of the fabric used in measures of pleasantness, unpleasantness and even arousal, suggesting that even after the induction of the RHI participants were able to discriminate between the two fabrics overall. Thus, the valence results of Experiment 1 may instead be explained by the emotional effects of the uncertainty of the seen tactile affectivity in our experiments. Uncertainty carries potentially unexpected consequences (Hsu et al., 2005): in Experiment 1 participants could not see the fabric touching the rubber hand, which could have led them to rate this mismatch between the felt affective certainty and seen uncertain tactile stimulation as less pleasant. By contrast, in the incongruent condition, both felt and seen touch had a degree of certainty. Taken together, these results suggest that although subjective embodiment is influenced by both affective certainty and congruency between felt and seen modalities, subjective pleasantness is influenced only by the former.

In conclusion, our results suggest that at the conscious experience of body-ownership participants more readily embody a rubber hand that ‘feels like’ their own hand, whereas they tend to resist the RHI when their own tactile affectivity does not match the vicariously perceived tactile affectivity on the rubber hand, even if the latter is more pleasant. We find that incongruences between the felt and seen tactile affectivity can disrupt the multisensory integration processes that lead to body ownership, over and above any effects of temporal synchrony, or felt valence. This is the first study to show that tactile affective congruency between direct and vicarious modalities modulates body awareness during multisensory integration, in ways similar to other amodal properties such as temporal or spatial congruency. We thus suggest that affective contingency in signals originating from different modalities may act as an important amodal precondition for multisensory integration, as predicted by developmental theories on the formation of the bodily self on the basis of affectively contingent, embodied interactions (Ciaunica & Fotopoulou, 2016; Fotopoulou & Tsakiris, 2017).

## Acknowledgments

This work was funded by a European Research Council (ERC) Starting Investigator Award for the project ‘The Bodily Self’ N313755 to A.F. Funding for the time of A.F. and L.C. has been partially provided by the Fund for Psychoanalytic Research through the American Psychoanalytic Association.

## Author Contributions Statement

All authors conceived and designed the experiment; M.L.F. performed the experiments and analysed data; M.L.F. and A.F. jointly wrote the manuscript; all authors read, corrected and approved the final manuscript.

